# Population structure and geographically structured reproductive strategies of the haplodiplontic seaweed *Dictyota dichotoma*

**DOI:** 10.1101/595587

**Authors:** Frédérique Steen, Verlaque Marc, Sofie D’hondt, Christophe Vieira, Olivier De Clerck

**Affiliations:** Phycology Research Group and Center for Molecular Phylogenetics and Evolution, Ghent University, Krijgslaan 281 (S8), Ghent B-9000, Belgium; Aix Marseille Universite, CNRS/INSU, Universite de Toulon, IRD, Mediterranean Institute of Oceanography (MIO), UM 110, GIS Posidonie, 13288 Marseille, France

## Abstract

Both mating system variation and the propensity of many seaweeds to reproduce both sexually and asexually, leave a strong imprint in the genetic structure of species. In this respect, we study the population genetic structure of *Dictyota dichotoma*, a common haplodiplont brown subtidal seaweed. This benthic species is widespread in the NE-Atlantic, from the Canary Islands and Mediterranean Sea to southern Norway, but lately populations have been reported from Argentina and South Africa. Phenology and reproduction of *D. dichotoma* was monitored year-round in four populations to investigate how the species has adapted to the steep thermal gradient in southern and northern ranges of its distribution. Thirteen microsatellites are developed in order to assess patterns of population diversity and structure across the biogeographic range, as shaped by past and present processes. Last, we assess the genetic structure of South African and South American populations and their relationship to the northern hemisphere populations.

Throughout its range, *D. dichotoma* shows a varying reproductive effort, with sexual reproduction being more abundant in the northern range. In contrast, the Mediterranean populations show a clear sporophyte dominance, suggesting that sexual reproduction is not the prime mode of reproduction, and indicating that the species potentially resorts to other modes of propagation as for instance fragmentation or apospory.

Genetic diversity is highest in the southern population decreasing gradually northward, indicative for a recolonization pattern after the demise of the last glacial maximum where these areas served as glacial refugia. European mainland populations show an isolation by distance pattern, while the population in the Canary Islands has its own genetic identity, being significantly diverged from the mainland population. Populations in South Africa and Argentina are seemingly introduced from mainland Europe, but no conclusion can be made on the exact timing of these introductions.

## Introduction

Natural populations are the ecological entity within which evolutionary processes are acting. Multiple evolutionary forces such as mutation, migration, selection and drift interact to shape the genetic structure of populations, and are heavily influenced by contingent historical, geographic and biological contexts (Loveless & Hamrick 1984). Glacial cycles and other global perturbations acting at geological time-scales are known to have dramatically influenced shifts in species distributions and spatial patterns of genetic variation within species (Hewitt 2000, Maggs *et al*. 2008). In addition to species-specific traits (e.g. intrinsic dispersal capacity), ecological factors affecting reproduction and dispersal, such as regional shoreline configurations, etc… are also particularly important in shaping connectivity at more local scales and hence influence genetic structure also. The environment can also select for different life-cycle strategies throughout a species range, with many taxa putting varying effort in sexual reproduction (Eckert 2002). While sexual reproduction produces new allelic combinations, asexual reproduction produces genetically identical offspring. The relative contribution of each strategy may thus lead to important differences in the genetic diversity and structure of populations across a species range (Eckert 2002, Balloux *et al*. 2003).

Macroalgae constitute good models to study the effects of mating system differentiation on population genetic structure, given their wide diversity of life cycles. Most macroalgae have biphasic life cycles with somatic growth in both haploid and diploid phases, potentially representing challenges for studies on their evolutionary ecology. Ideally, both life phases should be sampled, in order to explore connectivity between phases on larger geographical scales (Krueger-Hadfield & Hoban 2016). This can be particularly challenging as changes in habitat, fitness and disruption of the sexual cycle can lead to heavily biased ploidy ratios, giving rise to gametophyte (Van der Strate *et al*. 2002) or sporophyte dominances (Guillemin *et al*. 2008). Numerous algae have a capacity for both sexual and asexual reproduction (Santelices 1990), and often, asexual reproduction becomes more important towards geographical or ecological margins of species distributions (Eckert 2002, Billingham *et al*. 2003, Tatarenkov *et al*. 2005, Oppliger *et al*. 2014). High levels of asexual reproduction typically result in an increase in the number of repeated multilocus genotypes (ramets or clones), linkage disequilibrium and heterozygote excesses (Halkett *et al*. 2005), all of these in turn having a profound effect on the genetic composition and diversity of populations. In this respect, range-wide studies on the genetic structure of species that consider geographic variations in mating systems are of particular interest in a world of environmental change (Eckert *et al*. 2010).

To date, most large-scale studies dealing with patterns of population structure in brown algae have focused on species with heteromorphic life cycles as fucoids or kelp (Oppliger *et al*. 2014, but see: Couceiro *et al*. 2015). These studies have also several mechanisms of asexual reproduction as fragmentation in *Fucus radicans* and *Fucus vesiculosus* (Tatarenkov *et al*. 2005, Johannesson *et al*. 2011, Ardehed *et al*. 2015) or parthenogenesis in kelp (Oppliger *et al*. 2007, Oppliger *et al*. 2014). The greatest difference with *Dictyota dichotoma* lies within their heteromorphic life cycles. In fucoids gametophytes are reduced and integrated in the macroscopic sporophytes and in kelps the free-living gametophytes are greatly reduced in size as compared to the sporophyte generation. Reproductive characteristics, as mating system, frequency of recombination or also spatial or ecological separation of ploidy phases, greatly influences the genetic population structure of species. In this respect, the study of a haplodiplont organism with isomorphic ploidy phases such as *Dictyota dichotoma* is quite original and can expand current understanding on the drivers in range-wide population genetic structure of brown algae. *D. dichotoma* is a common species found in the subtidal zone of the northeast Atlantic, Mediterranean and Macaronesian coast. Recently, its presence has also been genetically confirmed for South Africa (Tronholm *et al*. 2010) and Argentina (Lopes-Filho *et al*. 2017), but their origins remained hitherto elusive. *D. dichotoma* is able to maintain appreciable biomass in the subtidal, even under high grazing pressures. This is attributed to the presence of grazer deterrent volatiles (Wiesemeier *et al*. 2007), and the ability to propagate by fragmentation. Reproduction occurs through an isomorphic alternation between dioecious gametophytes that produce male and female gametes and diploid sporophytes that produce tetraspores by meiosis. Fragmentation has been shown to be a mechanism of clonal propagation in *Dictyota* (Herren *et al*. 2006), and some species have also the ability to regenerate the sporophyte phase through the production of apomeiotic spores (Hwang *et al*. 2005). The latter has also been observed for *D. dichotoma* for a Mediterranean culture strain coming from Marseille (Steen, unpublished). Field observations suggest that gametophytes are generally lacking in the Mediterranean and therefore that sexual reproduction is rather an exception than the rule. Morphological observation however may not be adequate to draw inferences regarding the incidence of sexual reproduction, as gametophytes might be rare, cryptic or depending on specific environmental conditions and therefore be missed in the populations. As such genetic recombination via sexual reproduction, or the lack of recombination is better estimated using genetic data (Halkett *et al*. 2005).

The current geographical range of *D. dichotoma*, extending from Norway in northern Europe, as south as the Canary, has dramatically been influenced by temperature fluctuations across past glacial cycles. During Pleistocene glacial maxima, ice-sheets extended far downward central Europe, thereby making the coast uninhabitable for benthic organisms, only leaving southern regions viable for populations During interglacial, northern areas could be colonized again from these southern refugial populations (Hewitt 2000). At present, populations of *D. dichotoma* are subjected to large temperature gradients across this range. Temperature plays a prominent role in the control of sporogenesis and growth in the species (Bogaert *et al*. 2016), which could influence its reproductive strategy. As the relative contribution of sexual and asexual reproduction leaves a genetic signature within a species, we aim at integrating data from phenological observations with population structure of *D. dichotoma*. For this purpose we monitored the phenology and reproductive effort of *D. dichotoma* in the northern and southern stretches of its native range and relate these observations to its resulting genetic imprint. To do so, we developed co-dominant single locus microsatellites to explore how historical climatological events in conjunction with mating system differences have shaped the intraspecific genetic diversity of *D. dichotoma* across its range.

## Materials and Methods

### Phenology

Four populations (Carry-le-Rouet, Calanque de Sormiou, Wimereux, Goes, Table 2) of *Dictyota dichotoma* were monitored and sampled on a monthly basis from May 2012 - May 2013. If present, fifty individuals were sampled along a 50m transect, collecting 5 random individuals every 5 meter. The Wimereux population was confined to a series of interconnected shallow intertidal pools measuring approximately 10m². Ploidy was recorded whenever reproductive structures were visible. If not reproductive, thalli were scored as being sterile. A total of 221 out of 1437 samples were genotyped sampling haphazardly from the available dataset (67 in Goes, 53 in Carry-Le-Rouet, 47 in Calanque de Sormiou, 54 in Wimereux). Reproductive samples were used to confirm ploidy after genotyping: known haploids produced a single allele and known diploids produced either one or two alleles for homozygote and heterozygote loci respectively. Of the sterile individuals those exhibiting a single allele at all 13 loci were regarded as gametophytes. All samples were stored in silica and Carnoy’s solution for further examination. Mean monthly sea surface temperatures for each locality and at the time of sampling were extracted from NCEP Reynolds Optimally Interpolated Sea Surface Temperature data sets (Reynolds *et al*. 2002).

### Population genetic sampling, DNA extraction, microsatellite development and PCR reactions

A total of 18 populations of *D. dichotoma* was sampled (including the populations sampled for phenology), spanning most of the distribution of the species (Fig. 1). At each site, tissue samples from a minimum of 20 specimens separated at least 1m from one another were stored in silica gel.

**Figure 1.**
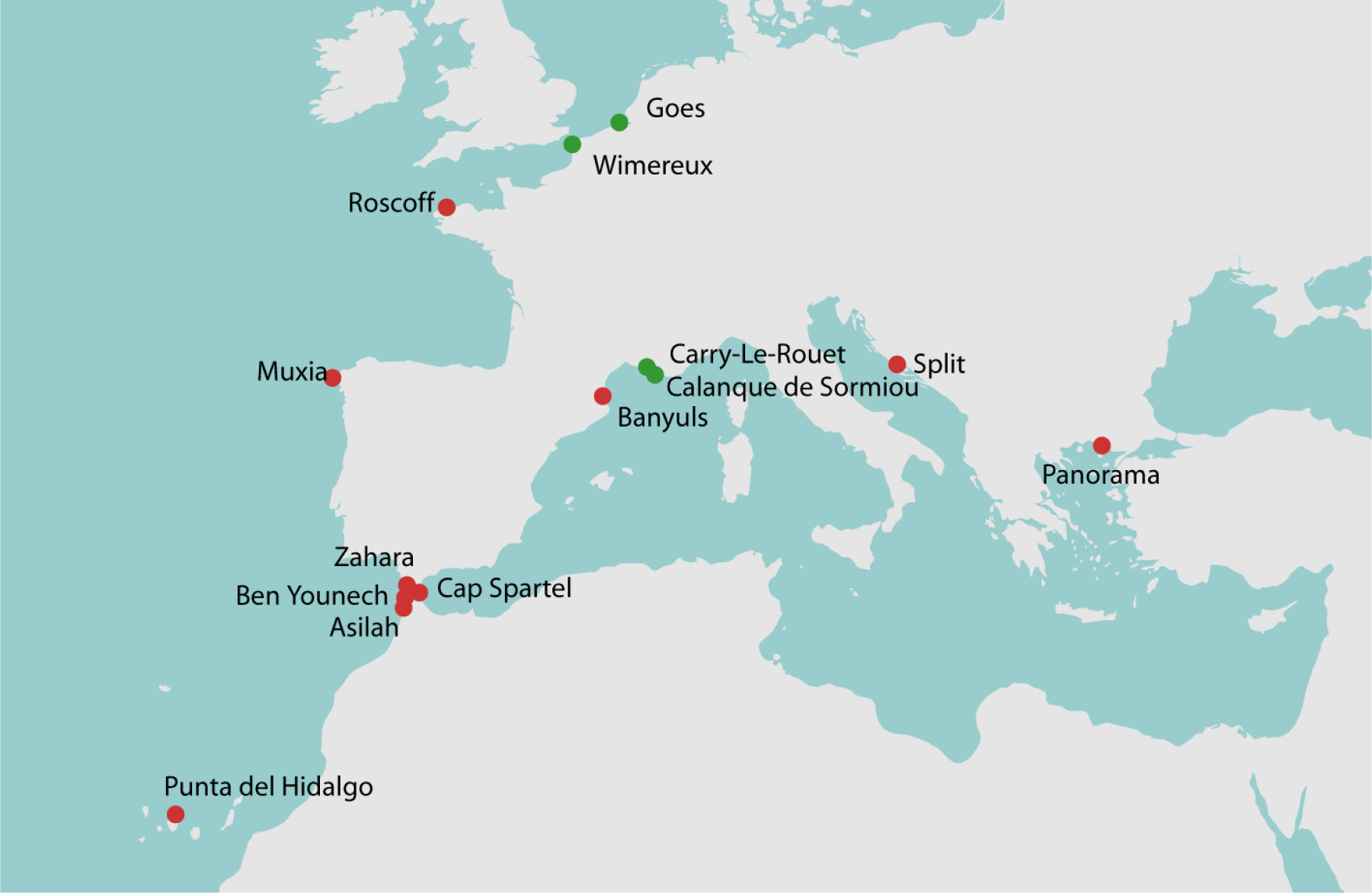
Sampling localities of *Dictyota dichotoma* in the NE Atlantic and the Mediterranean. Populations sampled in the phenological study are indicated in green.

Microsatellite development was based on genomic DNA extracted from one specimen using an adapted CTAB method (Steen *et al*. 2015) and sequenced with the Ion Torrent PGM sequencer. The run resulted in a total of 3⋅10^6^ raw reads which were analysed with the standard settings of QDD 3.1 (Meglecz *et al*. 2014), a pipeline implementing the Primer3 software (Untergasser *et al*. 2012) to design microsatellite primers. A total of 6391 microsatellite loci were obtained of which 55% di-nucleotide repeats, 17% tri-nucleotide repeats, 8% tetra-nucleotide repeats, 16% pentanucleotide and 4% hexa-nucleotide repeats. A set of forty di-to hexa-nucleotide repeats were selected according to the recommendations by Meglezc *et al*. (2014).

Thirteen microsatellites loci, estimated to be polymorphic based on a subset of samples, were amplified with fluorescently-labelled primers in 5 multiplex reactions as follows: DdA-DdC-DdH, DdB-DdB, Dd1-Dd14-Dd19, Dd3-Dd5-Dd10, Dd6-Dd7. PCR reactions were performed in a 12.5 µl reaction volume consisting of 1.25µl dNTPs, 1.25µl buffer solution, 0.625µl of each primer, 0.63 µl Taq polymerase, 0.5µl BSA, 0.5µl DNA and 6.25µl water. Annealing temperatures are as indicated in Table 1. The PCR program consisted of an initial denaturation of 96°C of 5 minutes, followed by 35 cycles of 1 minute each of 96°C, T_a_, 72°C and a final elongation of 72°C for 10 minutes. The allele size was scored and binned using the microsatellite plugin of Geneious ver. 7.1.7 (Kearse *et al*. 2012). Ploidy of individuals was estimated from genotypes as above.

**Table 1.**
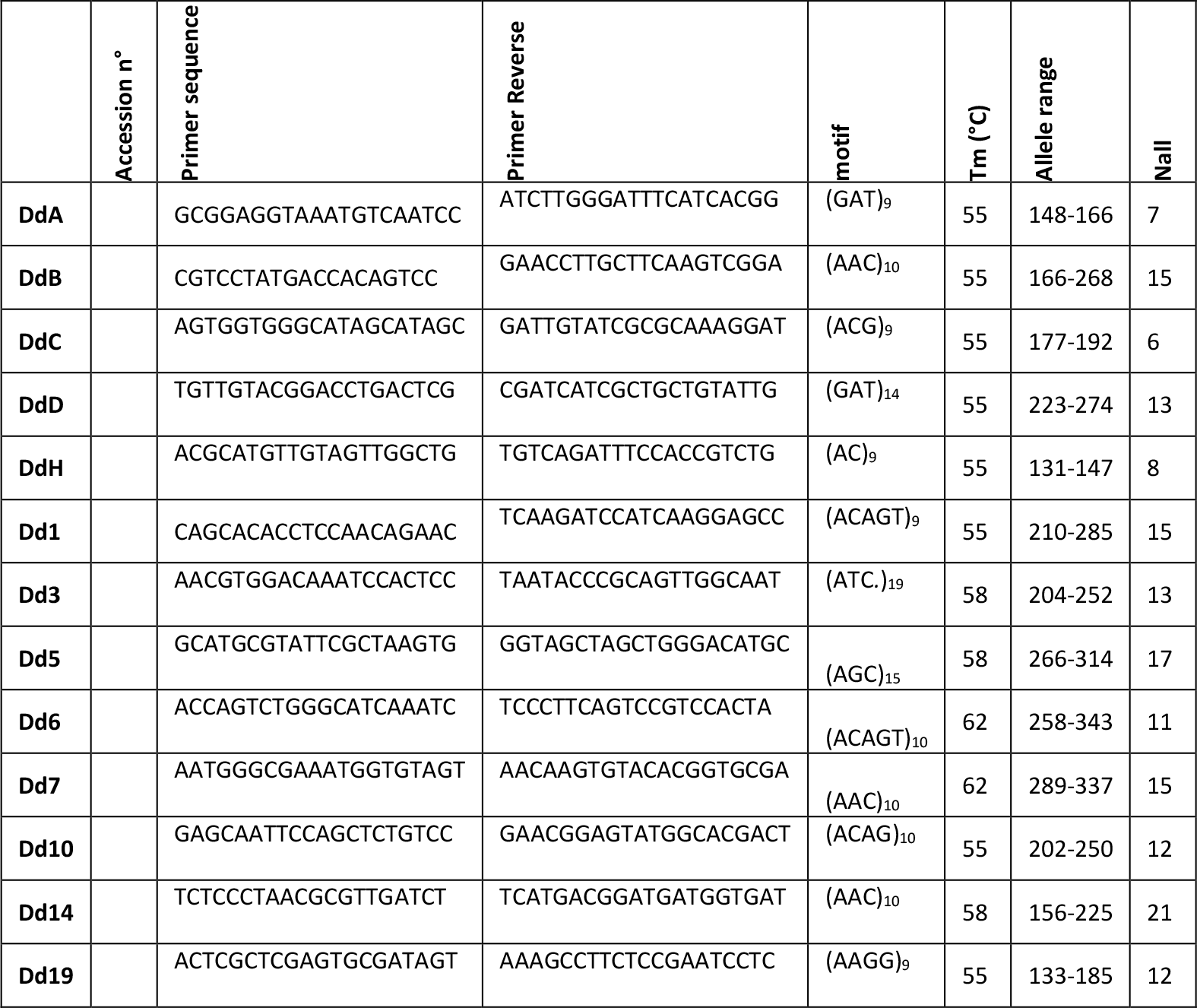
Microsatellite loci, GenBank accession numbers, forward and reverse primer sequences, motifs, melting temperatures, allele range and number of alleles (Nall)

### Population genetic analyses

#### Basic locus statistics and population genetic indices

The frequency of null alleles was directly estimated from the non-amplified samples of the haploid gametophytes, disregarding technical errors. The sporophyte data was analysed for null alleles, large allele dropout and stutter alleles using Micro-checker (Van Oosterhout *et al*. 2004). As the number of gametophyte samples was not large enough, diversity indices and subsequent analyses were calculated on the sporophyte subpopulation only. For the sporophytes the number of repeated multilocus microsatellite genotypes (MLGs) was computed using the function *mlg* in the *poppr* package (Kamvar *et al*. 2014) developed for R (R development core team, 2018). As repeated MLGs may occur due to asexual reproduction or repeated sampling of the same individual, for each MLG *Psex* was calculated as the probability that the MLG arose from a recombination event (Parks & Werth 1993). When *Psex* was greater than 0.05, the MLGs were considered being different genets and retained in subsequent analyses, else MLGs were removed in order not to bias estimates. Linkage disequilibrium (LD) between locus pairs was tested for each of the populations with Genepop (Rousset 2008) and p-values were adjusted with Bonferroni correction for multiple comparisons. Pairwise linkage disequilibrium is reported as the number of significant pairwise LD relative to the total number of pairwise comparisons in a population. Multilocus linkage disequilibrium of the sporophyte subpopulations was estimated for the total and clone-censored dataset as the standardized index of association (Agapow & Burt 2001) implemented in the package *poppr*. Linkage disequilibrium can also be caused by asexual reproduction, inbreeding, or by the presence of population structure. Significance testing was performed using 1000 permutations. Expected and observed heterozygosities were calculated with GENALEX version 6.5 (Peakall & Smouse 2012). HP-Rare (Kalinowski 2005) was used to compute the per locus mean expected number of alleles and the per locus mean expected number of private alleles, rarefied on the smallest sample size of 26 genes. FSTAT (Goudet 1995) was used to calculate F_is_ and to perform tests of HW-equilibrium. For each locus and over all loci F_is_ was calculated according to Weir & Cockerham (1984) and significance was tested by running 1000 permutations of alleles among individuals within samples. In addition to the full diploid dataset, F_is_ was also calculated for the clone-censored dataset as repeated MLG might bias estimates. Both G_st_ (Nei 1973) and Dest (Jost 2008) are calculated based on the clone censored dataset, employing the R-package *DEMETICS*. The measure for population differentiation G_st_ (Nei 1973) is highly dependent on within population genetic diversity thus this measure was complemented by calculating Dest (Jost 2008), as suggested by Meirmans & Hedrick (2011). The latter measure is more evident given the high mutation rate of microsatellites. Significance is based on p-values using 1000-fold bootstrap resampling. In order to correct for multiple comparisons, the Benjamini Hochberg procedure is used, with a false discovery rate of 0.05.

G tests (Goudet et al., 1996) in DEMEtics (Gerlach et al., 2010) used 1,000 bootstraps to provide 95% confidence intervals and p-values for global and per-locus estimates of Jost’s D (Jost, 2008). Benjamini & Yekutieli’s (2001) modified false discovery rate (FDR) method was chosen to adjust the significance threshold for pairwise FST p-values because it better controls Type I (α) error than the original FDR approach of Benjamini & Hochberg (1995) without the loss of power to distinguish meaningful genetic differentiation that occurs with the conservative Bonferroni correction (Narum, 2006)

#### Population structure in the native range

We employed a model free K-means clustering approach based on genetic distances to infer population genetic structure (Jombart *et al*. 2010), which does not assume unlinked markers and panmictic populations (Pritchard *et al*. 2000), and is therefore more convenient for species that reproduce clonally or partially clonally. In order to infer the number of genetic populations without prior information on population structure and sampling, we ran the function *find.clusters* in R package *adegenet* (Jombart 2008) on the complete diploid dataset. This function turns the original genotypic data into uncorrelated principal components and scores clustering solutions for different numbers of clusters using a Bayesian Information Criterion (BIC). The optimum number of clusters was anywhere between 15-20 (Suppl. Inf. 1) retaining all the PCs. We chose to proceed with a Discriminant Analysis of Principal Components (DAPC) using the sampling locality as a prior for the groups. In order to ensure the discriminant analysis (DA) with the population priors was not overfit by retaining too many PC’s, a preliminary full model was run including all PC’s. An optimal a-score was estimated in order to maximize the ability to assign individuals to clusters reliably. To do this, a permutation test was run with 1000 simulations, maximizing the a-score, by comparing the number of assignments to the “real” number of clusters to randomized clusters. The a-score measures the proportion of successful reassignments to the prior clusters or to random clusters, and is essentially a measure of ‘over-fitting’ of the model. The optimal number of principal components was then used to perform a final discriminant analysis in order to infer the distribution of genetic variance between sampling locations. As Wimereux was significantly differentiated from the other populations, obscuring the patterns between other populations (Suppl. Inf. 2), we chose to perform the DAPC on all of the native populations, but excluding Wimereux.

Correlation between genetic and geographic distances was tested for European populations (including or excluding Punta del Hidalgo) by performing an isolation by distance (IBD) analysis with the package *adegenet*. A Mantel test was performed to test the correlation between the linearized (x/1-x) Jost Dest pairwise estimates of genetic differentiation and the geographic distances among populations with the *mantel.randtest* function in R. Distances between localities were measured as the shortest path over the sea (oceanic distance). Probability values were obtained by 5000 random Monte Carlo resampling of the data using the *adegenet* package.

#### Potential origin of the introduced populations

To investigate the potential source population of the South-African and South-America samples, an additional DAPC was performed based on all sampled native populations (excluding Wimereux). Individuals from either South Africa (Kalk Bay and Strand) or Argentina (Villarino and Las Charas) were then subjected to the PCs of this DAPC, and individual samples were reassigned to potential source populations.

## Results

### Phenology

A total of 1425 specimen were sampled to assess the phenology of *Dictyota dichotoma* at 4 locations along the Atlantic and Mediterranean coasts, respectively (Table 2). Due to weather conditions and wave-exposure, the site Carry-Le-Rouet was not sampled in December and March. In Marseille, macroscopic thalli were virtually absent throughout July and August, when temperatures were highest (Fig. 2).

**Table 2.**
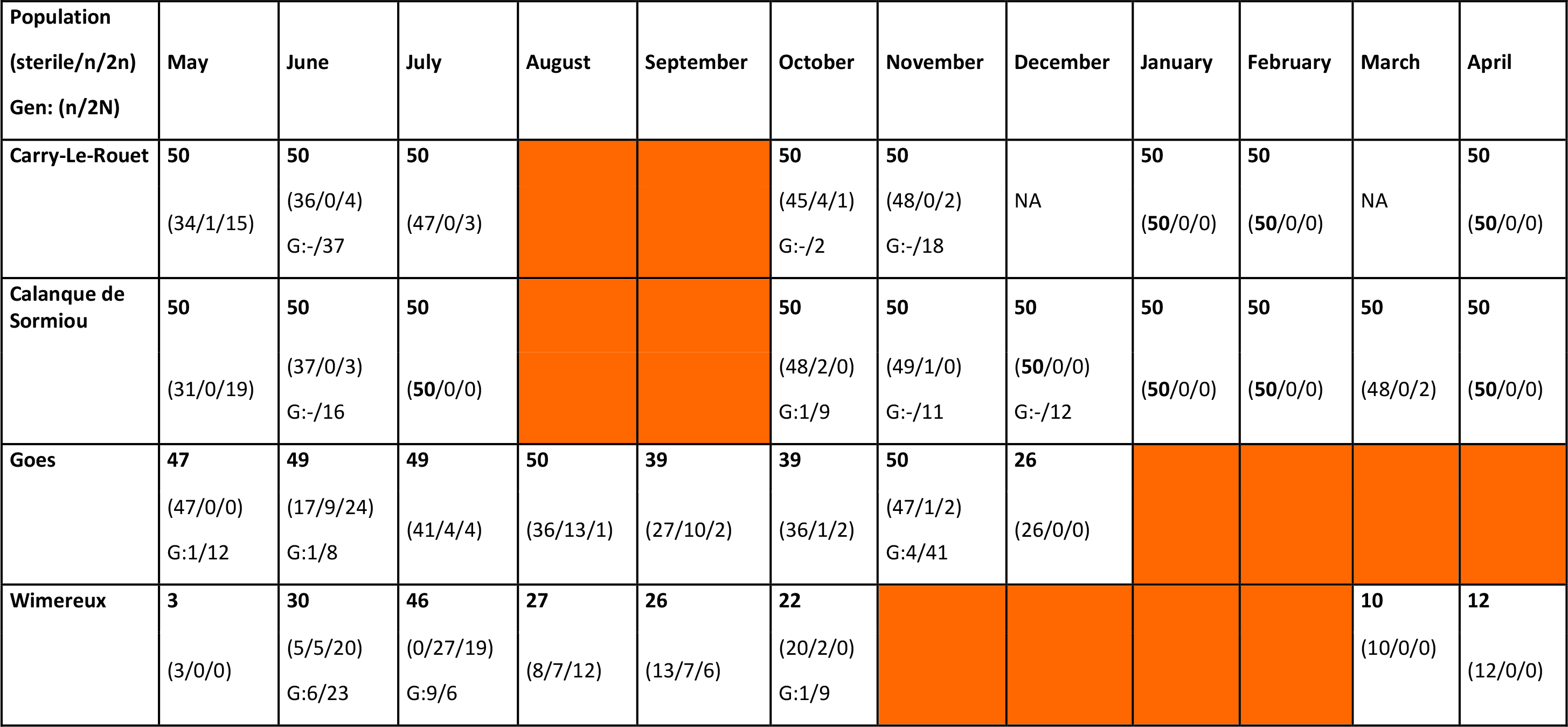
Localities sampled for phenology. Total numbers of samples are indicated in bold, numbers between brackets indicate the number of morphologically observed fertility (sterile/gametophyte/sporophytes) respectively. When genotyped, inferred ploidy is indicated as G: # haploid/# diploid. Months when conditions did not allow to sample are indicated by NA. If no thalli were present, months are indicated in orange.

**Figure 2.**
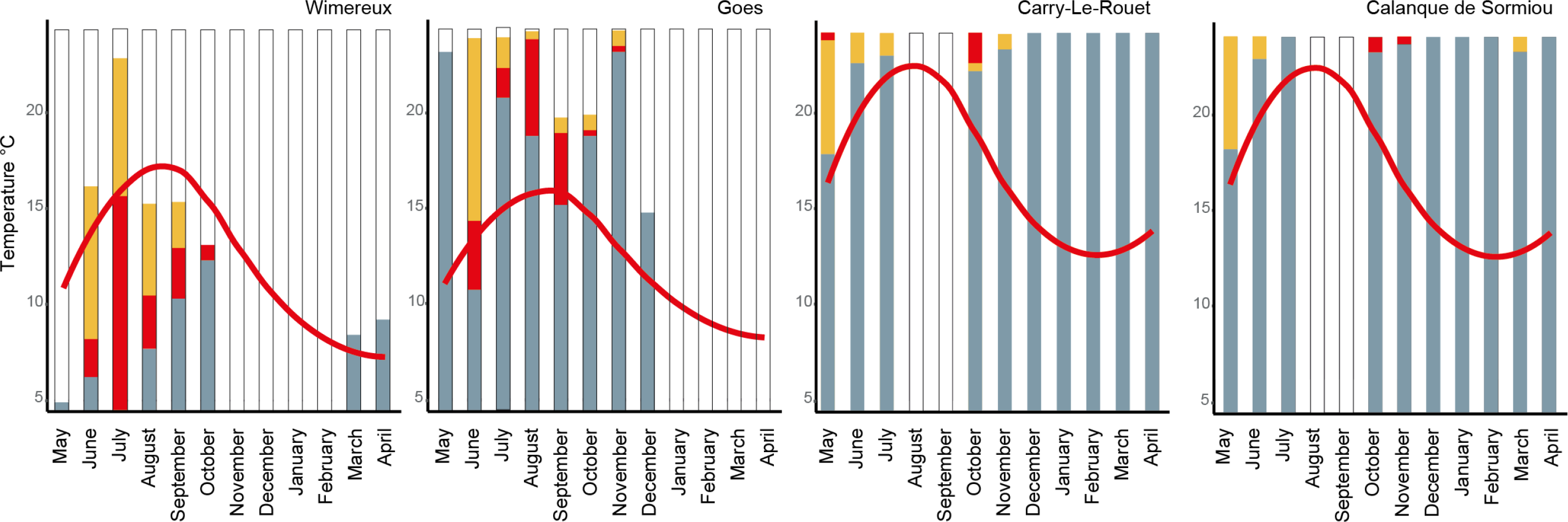
Relative abundance of sterile (grey), gametophyte (red) and sporophyte (yellow) samples per month. The height of the bar indicates 50 samples, the white proportions are thus the non-sampled species given the low abdunance of *D. dichotoma* at this time of the year. The red line represents the local mean monthly sea surface temperatures.

*D. dichotoma* was absent in Goes from January to April and in Wimereux from November till February, when temperatures were lowest. Out of 900 specimens sampled in both two sites near Marseille, fertile gametophytes were extremely rare and mostly observed in autumn: 1 male and 3 female gametophytes in October in Carry-Le-Rouet, and 2 male and 1 female in October and November for the Calanque de Sormiou population. In spring only one fertile male was observed in May in Carry-Le-Rouet. In contrast, the populations in Wimereux and Goes show fertile gametophytes mainly from June to September, and in lower frequencies extending throughout October and November respectively. Fertile porophytes were present from June till September for Wimereux and from June till November in Goes. In Marseille the proportion of fertile individuals was much lower than along the Atlantic coast. Almost all individuals in Wimereux and the majority of the individuals in Goes were fertile in June and July.

### Population genetic analyses

#### Basic locus statistics and population genetic indices

For the diploid samples, no evidence was present for stutter alleles or large allele dropout. A general homozygote excess is noted for the diploid samples when comparing observed and expected heterozygosities (Table 3). This is not caused by null alleles, given the very low proportion of non-amplification (maximum 1% for Dd 1 and Dd7) of known haploid samples. The total missing data for the diploid samples was 1%, but this appears to be an artefact of the multiplex PCR, rather than null alleles. Therefore, we did not consider null alleles to have an effect on further analyses. After removal of individuals having greater than 10% missing data (i.e. more than 1 locus not amplified out of 13) the diploid dataset totalled to 458 individuals with 0.2% missing data. Missing data per locus did not exceed 0.9% (Locus DdD and Dd7) and missing data per population did not exceed 1.0% (Villarino).

**Table 3.**
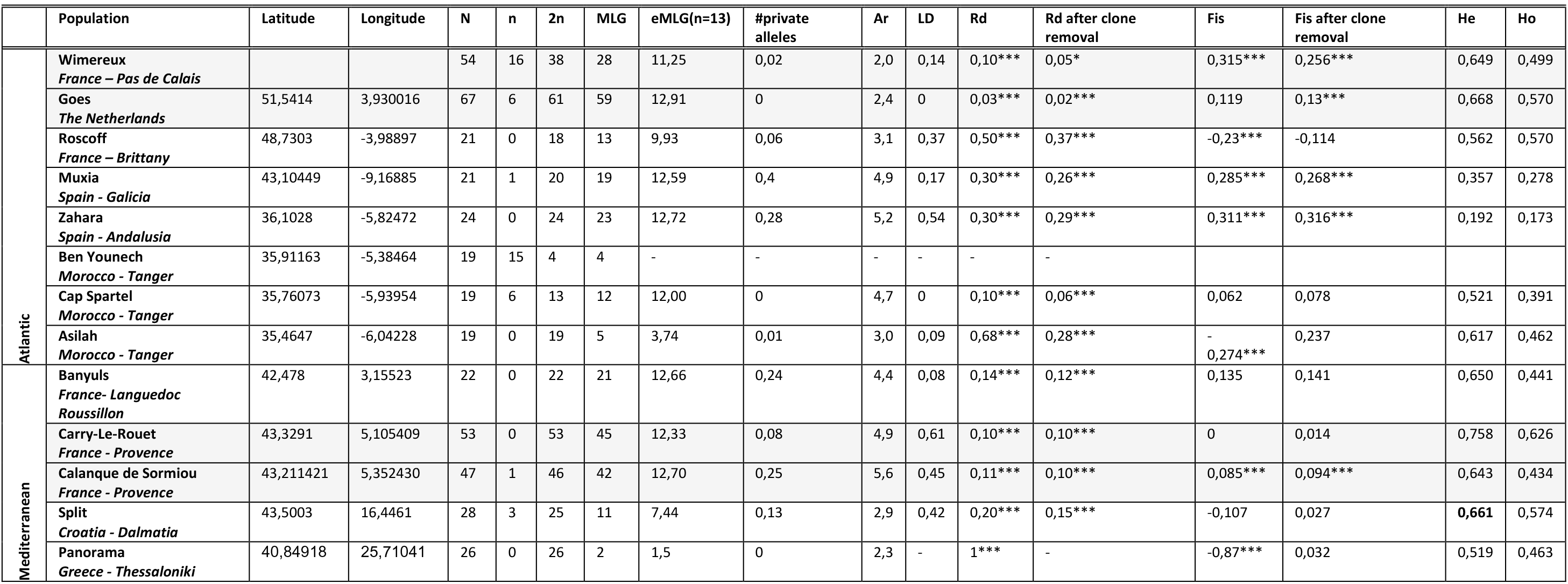
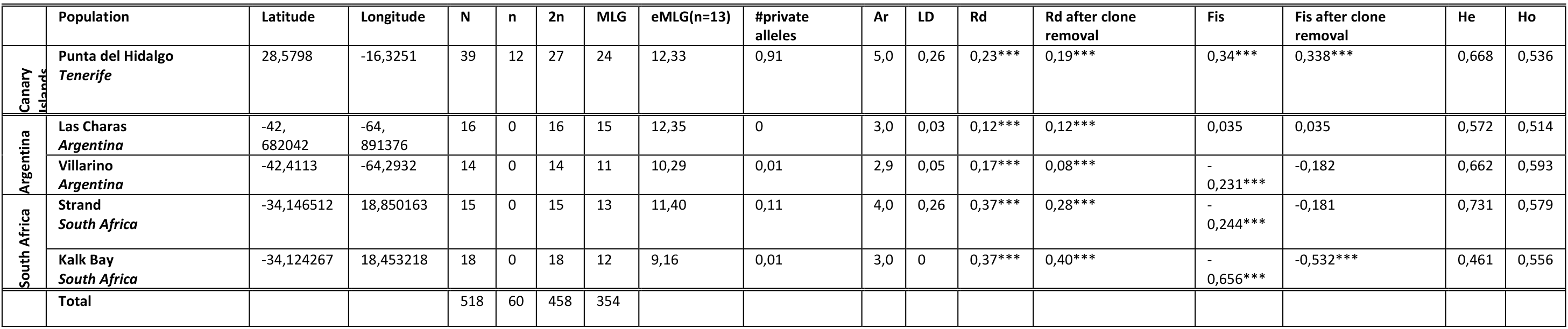
Population specific genetic diversity measures. Populations in grey are also studied for phenology in this study. Following measures are reported: N (number of samples), n (gametophytes), 2n (sporophytes), MLG (number of multilocus genotypes), eMLG (standardized number of MLG rarified to 13 diploid samples), #private alleles (mean number of private alleles per locus, rarified to 13 diploid samples), Ar (mean allelic richness, rarified over 13 diploid samples), LD (proportion of significant linkage disequilibrium between pairs of loci), Rd (multilocus linkage disequilibrium), Fis (inbreeding statistic), He (expected heterozygosity, observed heterozygosity). Fis and Rd were also calculated after clone censoring. Significance level is indicated as * for p<0.05, ** for p <0.01 and *** for p<0.001

All MLG had a probability of *Psex* lower than 0.05, except for two MLGs from Wimereux. Both copies of the respective MLGs were retained within the clone corrected dataset (MLG 133 (W17-6, W19-6) & MLG135 (W7-10, W26-6)) in order to calculate F_is_ for the clone-censored dataset. Pairwise linkage disequilibrium (LD) within population was highest in the Marseille populations, together with Split and Zahara. Multilocus LD was significant in any of the populations (Table 3). Values were highest at the southern and eastern edge of the native range (Panorama and Asilah respectively) and in the South African populations, as well as in Zahara, Muxia and Roscoff. Clone-censoring did not reduce these values greatly.

#### Genetic and genotypic diversity

Genetic diversity indices for the sporophytes are reported in Table 3. The rarefied number of MLGs on a total of 13 individuals was lowest for Panorama (1,5), Asilah (3,74), Split (7,44) and for Kalk Bay (9,16). Allelic richness is highest in the southern populations (Calanque de Sormiou, Zahara, Punta del Hidalgo, Muxia, Cap Spartel) and declines towards the edges of the native distribution of the species (in the north: Wimereux and Goes and in the east: Split and Panorama). Argentinian populations (Villarino and Las Charas), together with Kalk Bay show a comparatively low allelic richness. Strand in South Africa has an allelic richness similar to the southern European populations. High allelic richness is observed in populations with highest mean number of private alleles per locus (Zahara, Muxia, Banyuls and Calanque de Sormiou). The highest mean standardized (n=13) number of private alleles (0.91) was noted for Punta del Hidalgo. In the introduced populations of Kalk Bay, Las Charas, and Villarino, almost no private alleles were observed, as in the northern populations Goes and Wimereux and the edge populations of Panorama and Asilah. A heterozygote excess (negative F_is_) was noted for Roscoff, Asilah, Panorama, Strand, Kalk Bay and Villarino. After clone-censoring, a significant departure from HW-equilibrium was only noted for Kalk Bay representing heterozygote excesses. Significant homozygote excesses were apparent for Wimereux, Goes, Muxia, Zahara, Calanque de Sormiou and Punta del Hidalgo.

#### Population structure, population differentiation and isolation by distance

After 1000 simulations, the optimal number of principal components was 14 (Suppl. inf. 1). This number of principal components retained 68% of the total variance. The final model using 14 principal components revealed overlap between individuals from different sampling localities. This model was also able to reassign 75% of the individuals back to the original sample. The a-score optimization shows that the 14 retained principal components give a high reassignment success, indicating that sampling locality has an important effect on the clustering of this data. The scatter plot (Fig. 3a) visualizing the between-group structure of the native populations excluding Wimereux (but see Suppl. Fig 2), reflects the geographical continuum along the Atlantic and Mediterranean coast on the first axis. Along the second axis, the Canary island population is separated from the mainland samples. Pairwise differentiation of populations as measured with Fst and Dest are summarized in the supplementary (Suppl. Fig. 4). Globally, the most differentiated populations are the Canary island population Punta del Hidalgo as well as Wimereux and Goes depending on the indices used. Geographically-close population pairs could be highly (Wimereux/Goes) or weakly (Asilah/Cap Spartel) differentiated, depending on the region considered. A pattern of isolation by distance (IBD) was observed for the European populations (p<0,05; r²=0,45) (Suppl. Fig. 3), although stronger when excluding the Canary Island population Punta del Hidalgo (p<0,01; r²=0,58) (Fig. 4).

**Figure 3a.**
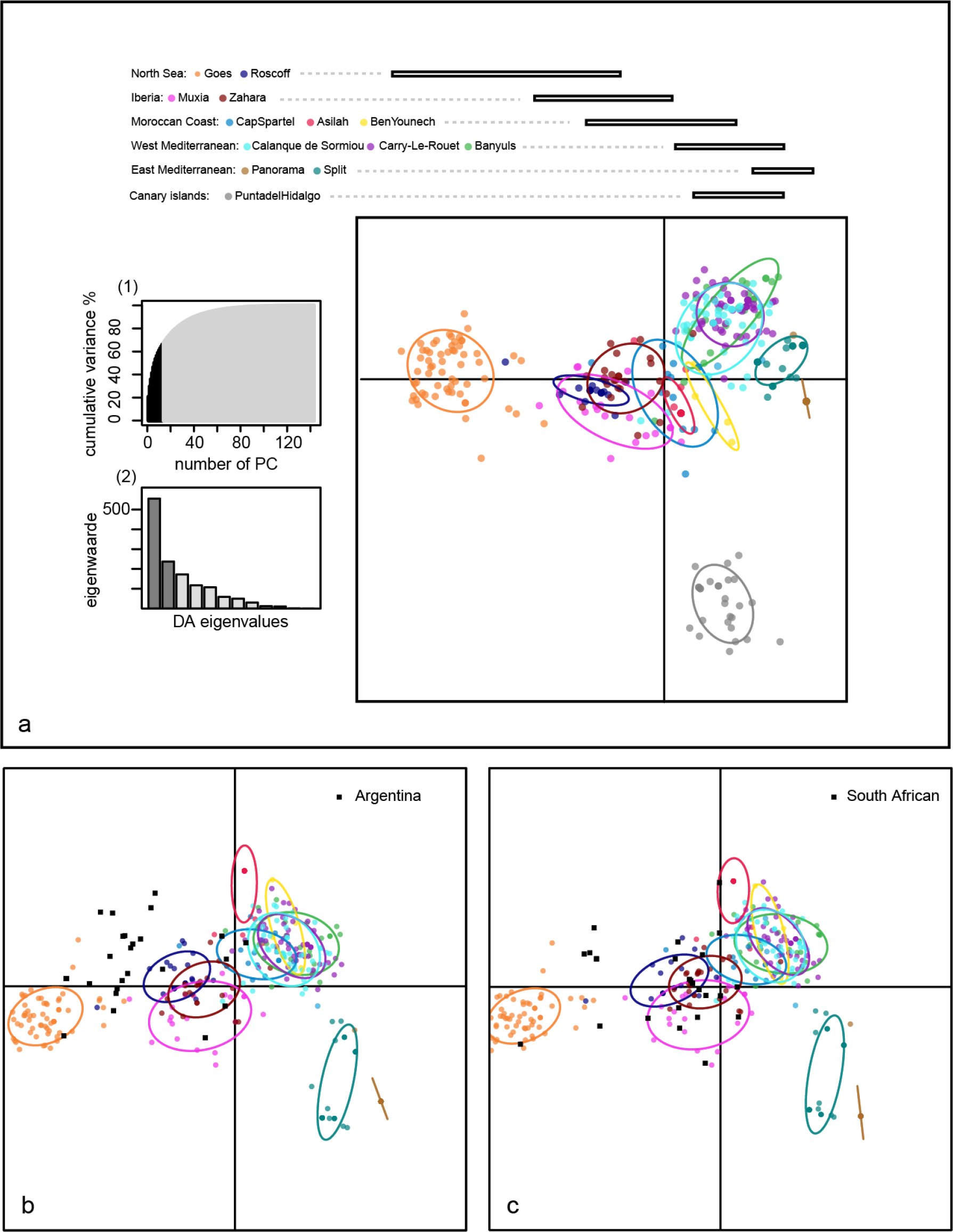
Scatter plot of the Discriminant Analysis of principal components. The individuals plotted along the first and second axis are colored according to sampling location as indicated in the legend. To clarify the geographical origin of the different samples and the geographical pattern of the scatter plot, individuals are post hoc grouped into regions (bars above). The bars above the plot represent the repartition of these geographical groups along the first axis. Cumulative variance explained by the eigenvalues of the PCA is represented in inset (1). Inset (2) displays a barplot with eigenvalues for the retained discriminant functions. Represented discriminant functions are in dark grey (b) reassigned Argentinian samples are indicated in black. c) reassigned South African samples are indicated in black.

**Figure 4.**
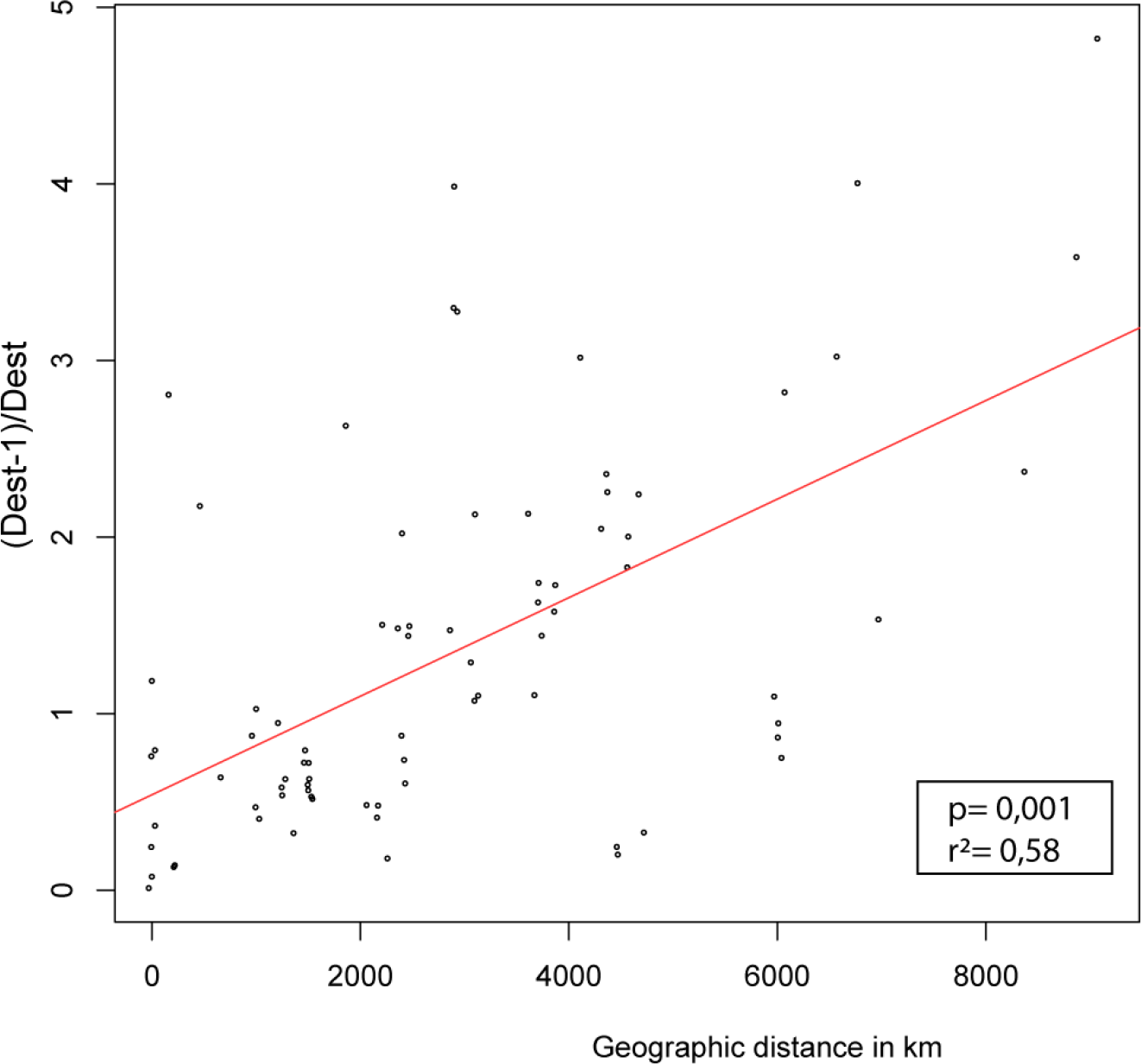
Isolation by distance in *D. dichotoma*, with. Linearized Jost Dest as genetic distance and shoreline distances, with r² the correlation coefficient of the regression line and p-value indicating significance of the Mantel-test. The population of Punta del Hidalgo is not included.

#### Potential origin of introduced populations

In order to trace genetically introduced populations to their putative region of origin, we reassigned individuals from either Argentina or South Africa onto the training dataset. South African individuals were plotted widespread over the European scatterplot (Fig. 3c). Individuals were mainly reassigned to Zahara (n=15) and Goes (n=8), but also to Muxia (n=3) Split (n=2), and 1 individual each to Calanque de Sormiou, Cap Spartel, Banyuls and Asilah. Samples from Argentina seem to be genetically more similar (Fig. 3b). Individuals are reassigned to Goes (n=13), Muxia (n=8), Asilah (n=5) and Cap Spartel (n=4).

## Discussion

### Phenology and geographical shifts in reproductive strategies

The growth and reproductive season of *D. dichotoma* is variable throughout its latitudinal range, suggesting that it is regulated by temperature. In the Marseille area populations are present throughout most of the year and absent in August and September. Mediterranean populations are thus largely concurrent with the phenology in the Canary Island populations as described by Tronholm (2008), with thalli disappearing the few months when SST higher or equal to 24°C (Canary islands: August= 24,0°C, Marseille: July= 23,8°C). In contrast, in the northern range macroscopic thalli of *D. dichotoma* were absent mostly during winter when SST were lowest. Increasing temperatures in spring promote sporophyte growth followed by sporogenesis. Logically, only after this stage gametophytes are observed. Richardson (1979) noted for *D. menstrualis* (Hoyt), Schnetter, Hörnig et Weber-Peukert (as *D. dichotoma*) that zygotes need summer conditions to germinate. If germlings were allowed to grow until a 7-9 cell stage, they were able to survive 21 weeks of winter conditions. For *D. dichotoma* experiments have indeed shown that sporophyte germlings from Roscoff can be conserved with limited growth at temperatures as low as 8 °C (Bogaert *et al*. 2016). These low temperatures also inhibit sporogenesis. These findings together with dominance of sporophytes in early spring could be an indication that *D. dichotoma* hibernates as small sporophyte germlings.

The highest fertility rates were first observed in spring for the northern (June SST 12.9 °C for Goes and July SST 13.5°C for Wimereux) and later on for the Mediterranean populations (May SST 16.4°C). Canary Island populations were most fertile at the end of winter (February SST 19.4°C and March SST 19.0°C). Roughly, fertility was observed when SST measure between 13° to 23° C suggesting that temperature regulates sexual reproduction. Given the discrepancy of the timing of observation of fertile individuals between different latitudinal regions, we speculate that temperature rather than photoperiodism plays the most significant role in the onset of sexual reproduction of *D. dichotoma*.

Overall, a general dominance of sporophytes over gametophytes was observed, both at the onset and demise of the growing season, in northern as well as in southern regions. Over all genotyped samples we found 60 haploids on a total of 533 individuals suggesting a gametophyte: sporophyte ratio (1 : 8,8) greatly deviating from the proposed theoretical √2:1 equilibrium ratio for biphasic and dioecious seaweeds that obligately cycle between the two ecologically equivalent phases, in the absence of apomixis (Destombe *et al*. 1989, Thornber & Gaines 2004). A biased sporophyte to gametophyte ratio was even more prominent for the Mediterranean populations, where over a total of 900 samples only 5 fertile gametophytes were observed as opposed to 49 fertile sporophytes. This corroborates earlier suggestions that dictyotalean species reproduce predominantly the sporophyte phase, and that sexual reproduction only happens occasionally (Williams 1905, Foster *et al*. 1972, Allender 1977, King & Farrant 1987, Phillips 1988, Mayhoub & Billard 1991, Hwang *et al*. 2005). This also indicates the possible wide-spread occurrence of apomixis for the species. Sporophytic dominance in *Dictyota* has been related to several mechanisms as direct development of apomeiotic tetraspores into new sporophytic thalli (apospory), greater longevity and vegetative reproduction of the sporophyte generations (Phillips 1988). Our data cannot distinguish between these mechanisms, but observations of recycling of the sporophyte phase in culture strains of the Marseille populations, indeed suggest a possible role for apospory. Also, the low fertility rate in combination with the abundance of *Dictyota* in the populations of Marseille, suggests that asexual propagation by fragmentation also contributes to the deviation from the theoretical √2:1 equilibrium ratio. This sporophyte dominance however, is not exceptional, and has been noted for many species of algae (De Wreede & Klinger 1988). We do cleary observe a southward decline in th investment of sexual reproduction, suggesting that environmental differences induce this gradient.

### Genetic signatures of varying mating systems

Differences in reproductive effort highly influence the genetic structure in populations, as only sexual reproduction is able to create new allelic combinations through recombination, while asexual reproduction results in identical genotypes. This leads to strong genetic imprints in the population structure of organisms reproducing clonally or partially clonally. In Panorama (Greece), and to a lesser extent in Split (Croatia) and Asilah (Morocco), we observe key signatures of clonal reproduction, as shown by repeated multilocus genotypes, linkage disequilibrium and a significant heterozygote excess (Halkett *et al*. 2005). In fact, in all of the sampled populations we do observe repeated multilocus genotypes and linkage between the markers, although inbreeding statistics do not indicate a significant heterozygote excess, and in some cases even homozygote excess. For the monitored populations in northern Europe sexual reproduction as a haplodiplont alternation of generations is observed (Wimereux, Goes), but also in Argentina (Gauna *et al*. 2013). In other Mediterranean populations the gametophyte phase was barely observed, both based on morphological examination and genetic data, corroborating that a high proportion of the population does not originate from sexual reproduction alone. These patterns can be caused by a reproductive system where clonality or asexual reproduction play a role, thereby disrupting equilibrium. High negative values of Fis are the ultimate signature for asexual reproduction (Halkett *et al*. 2005), but Fis vary greatly in systems with partial clonality. In addition, both positive and negative Fis seem possible and are observed for D. *dichotoma*. Negative values alone can’t be interpreted as evidence for asexual reproduction, but could be caused by intermediate rates of clonality in finite population (Reichel *et al*. 2016). Unfortunately, as recommended for biphasic organisms (Krueger-Hadfield & Hoban 2016), it is difficult for D. *dichotoma* to sample haploids at sufficiently large numbers, compromising the analysis of genetic differentiation between the gametophyte and sporophyte generations. Significant differentiation between both phases would be an extra indication of asexual reproduction, as gene flow between phases would be restricted leading to differentiation (Coyer *et al*. 1994, Wattier *et al*. 1997). However, the observed lack of gametophytes is an indication that the species resorts to other modes of reproduction.

### Population structure and connectivity

Seaweeds with limited dispersal capacities can show a pattern of isolation by distance (IBD), with populations in geographic proximity being genetically more similar than those separated by larger geographical distances. When genetic drift is greater than gene flow, this pattern between geographic and genetic distance quickly erodes (Wares 2002), and at least for intertidal fucoids and for some kelps, signals of isolation by distance quickly fade at larger distances (Fraser *et al*. 2010, Olsen *et al*. 2010, Neiva *et al*. 2012). This pattern of increased isolation by distance has been refuted for many fucoid algae and kelps. Often sharp genetic breaks are observed which are not correlated with distance (e.g. Tellier *et al*. 2009, Macaya & Zuccarello 2010, Neiva *et al*. 2012). For the subtidal *D. dichotoma* we do clearly observe this IBD pattern along the whole of the northeast Atlantic and Mediterranean coast, suggesting a stepping stone model of gene flow between European populations. We did not observe any sharp genetic breaks over the Atlanto-Mediterranean transition as suggested for different marine organisms (Patarnello *et al*. 2007), suggesting that populations of *D. dichotoma* are connected and able to exchange propagules contributing to the genetic pool of neighbouring populations. Even though including the Canary Island population in the analyses does not erases this signal, the latter seems to be even more diverged, with divergence measures consistently higher for any given distance. The stretch of ocean between the mainland and the Canary Islands thus may represent a barrier for genetic exchange, and contribute to the divergence of this population from the mainland. The absence of *D. dichotoma* further south along the Moroccan coast indicates this as a plausible hypothesis. Analogous to Punta del Hidalgo, Wimereux in Nord-Pas-de-Calais is also highly distinct from other European populations, based on measures of genetic differentiation. But in contrast, this population shows a homozygote excess, low allelic richness and a high proportion of fixed loci (5 out of 13), all of which are indicative of inbreeding. As this population was sampled from a tidal flat, isolated on a sandy stretch of coast, our results suggest that this sampling location is merely a small, isolated population with little potential for genetic exchange. Taking into account the small size and the secludedness of this population, genetic drift could have caused fast differentiation from the other populations (Suppl. Fig. 2).

### Interplay between historical range dynamics and geographical parthenogenesis

A pattern of southern richness and northern homogeneity (Hewitt, 2000) is evident in *Dictyota dichotoma*. Southern populations have a high proportion of endemic alleles and higher genetic diversity. This is clearest in the populations of Punta del Hidalgo on the Canary Islands, the Iberian peninsula (Muxia and Zahara), and to a lesser extent in the western Mediterranean (Banyuls and Calanque de Sormiou). These areas have previously been designated as potential glacial refugia for seaweeds (Van den Hoek *et al*. 1990, Hoarau *et al*. 2007, Maggs *et al*. 2008), and it may well be the case they acted also as persistence areas where *D. dichotoma* has survived during past climatic cycles. A gradual decline of genetic diversity and lack of private alleles in northern populations support a scenario of northward colonization of these previously glaciated areas after the demise of the last glacial maximum (LGM, ca. 20,000 ya). As the Canary Island population Punta del Hidalgo is highly differentiated from the population on the mainland coast, this indicates a minimal contribution to the genetic diversity of this area. Expansion after the last glacial maximum to the north, therefore most probably came from the Iberian or Western Mediterranean refugium.

The influence of Pleistocene ice ages is correlated with geographical parthenogenesis in the case of *D. dichotoma*, an association that is often observed (Kearney 2005). Reproductive mode of *D. dichotoma* seems highly dependent on the location in its native range. Sexual reproduction declines towards the edge populations in the eastern Mediterranean. In northern populations, formerly glaciated, sexual reproduction is readily observed in conjunction with low genetic diversity caused by recolonization. In contrast, populations within the eastern Mediterranean show evidence for high amounts of asexual reproduction together with low genetic diversity. In this respect, they could be considered rear edge populations in marginal habitats, maladapted to prevailing ecological conditions. These trailing edge populations are often located in localities where suboptimal conditions prevail, in our case assumed to be elevated SST. Small population size and low gene flow are possibly resulting in increased clonal reproduction (Billingham *et al*. 2003, Arnaud-Haond *et al*. 2006, Beatty *et al*. 2008). In the end, this clonal reproduction could lead to the loss of genetic diversity and adaptive potential. The western Mediterranean Sea harbours a high genetic diversity reflecting the southern richness signal. Nevertheless, sexual reproduction seems not to be the rule as indicated by the deficit of fertile gametophytes and high linkage disequilibrium in the Marseille populations. This could be the first sign of the onset of genetic erosion of these genetically diverse populations. The low rates of fertile thalli are also indicative of the western Mediterranean representing a suboptimal habitat today, and that current-day persistence is facilitated by the mechanism of asexual reproduction. Asexual reproduction could maintain the presence of a species in a marginal habitat, by keeping locally co-adapted gene complexes, or by by-passing unachieved reproductive tipping points. In the case of *D. dichotoma*, it is clear that the observed genetic structure is caused by the interplay between historical range dynamics and the geographical parthenogenesis.

### A non-native origin of Dictyota in the southern hemisphere

The southern hemisphere populations of Argentina and South Africa are genetically related to the mainland European populations, although current data makes it difficult to narrow down the putative source populations. Private alleles are almost lacking from both of Argentina and South Africa suggesting these introductions are recent and at least not predating a continental European – Canary Island divergence. Individuals from Argentina are reassigned to a subset of populations to which South African samples are reassigned. It is also clear that South Africa harbours a greater genetic diversity than Argentinian samples. Therefore we do not exclude the possibility that *D. dichotoma* from Argentina is introduced secondarily from the South-African populations. Alternatively, these introductions could be independent events. The greater genetic diversity in South-Africa (with samples showing affinities to both Mediterranean and NE-Atlantic populations) could also be explained by independent introductions from different European sources. High allelic diversity and linkage in Strand and Kalk Bay is consistent with this hypothesis and could point to multiple introduction events. Exact origin of southern hemisphere populations however remains speculative, and should be addressed with a more thorough population sampling of both native and invasive range. The study of these populations with a high number of SNPs could enable revealing the fine scale introduction history of this species, allowing us to also estimate the timing of this event.

## Conclusion

Climatological cycling in the northeast Atlantic, has clearly contributed the genetic structure of *Dictyota dichotoma* populations. A northward decline in genetic diversity is clearly observed. In contrast, we do observe an inverse gradient of reproductive effort, with increasing evidence for asexual reproduction towards the eastern Mediterranean. Although high levels of genetic diversity and private alleles are found in the Western Mediterranean, there are indications that sexual reproduction is scarce thereby showing that the area under present day climatological conditions is not a stable rear edge and that it has turned into a marginal habitat. *D. dichotoma* is very plastic, adapting its phenology and reproductive strategies throughout its wide and environmentally diverse range, with variation in patterns of genetic diversity among populations.

## Appendix

**Supplementary 1:**
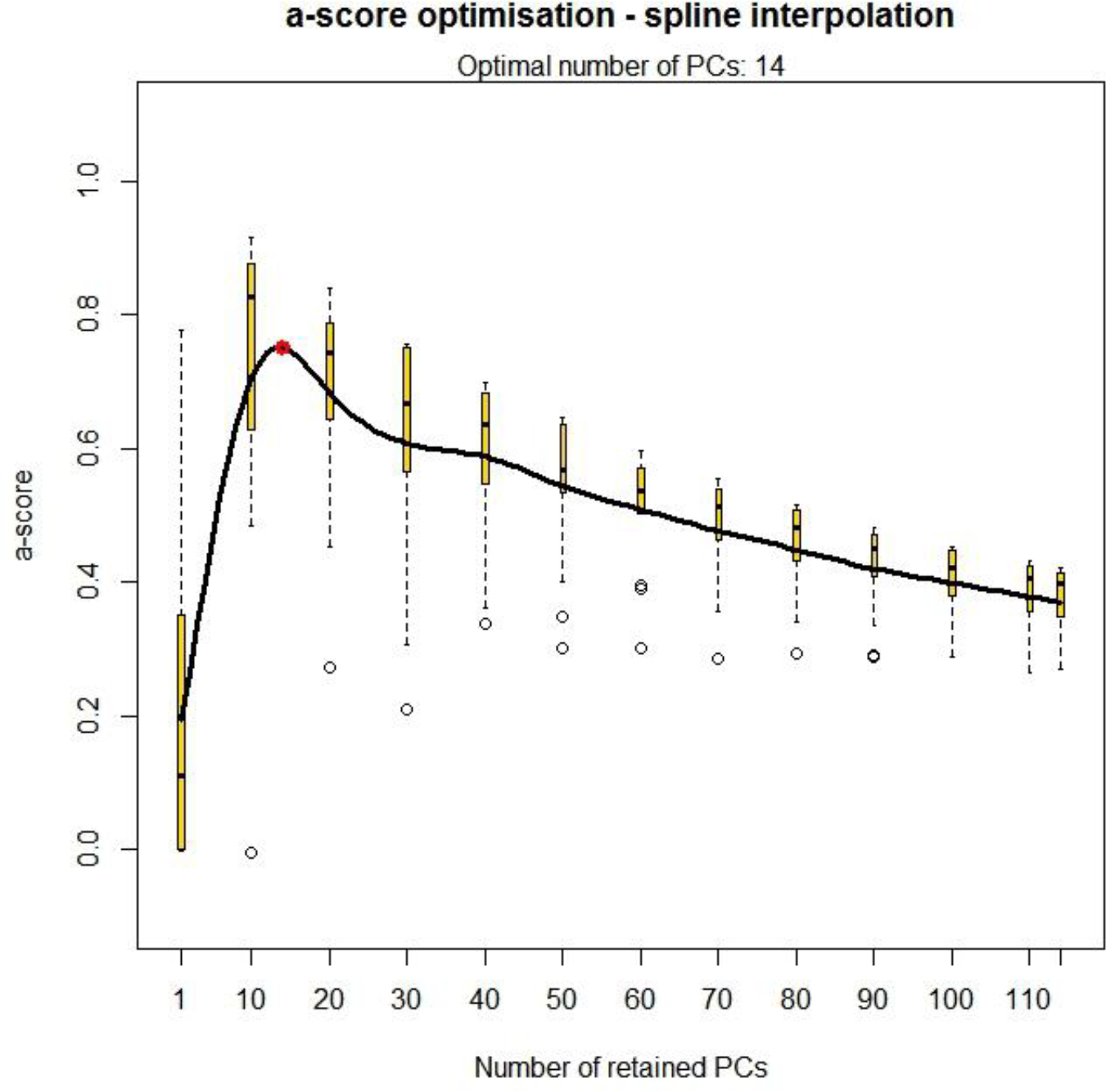
Graph representing the optimal number of retained PCs.

**Supplementary 2.**
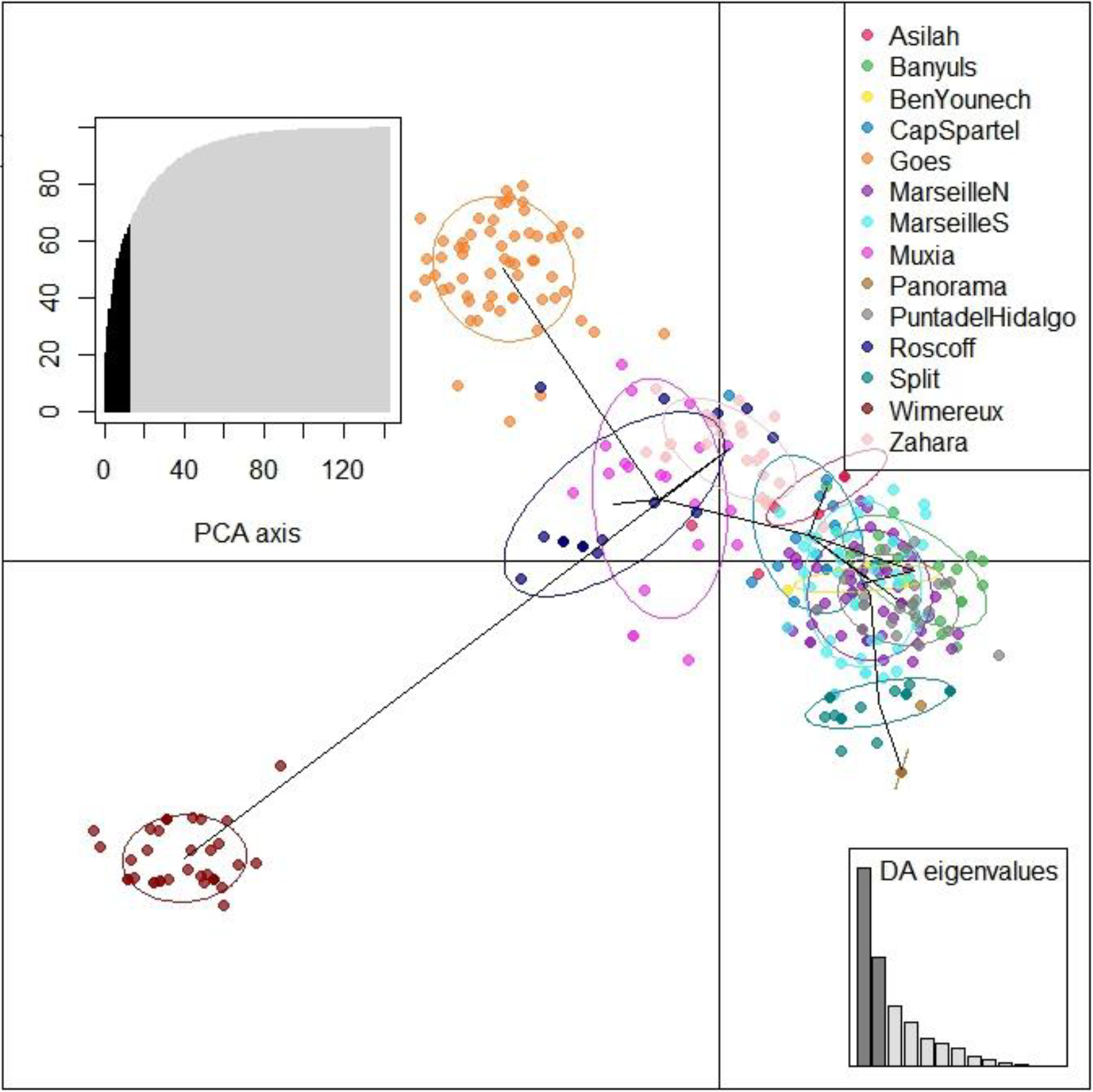
DAPC for all native populations including Wimereux, retaining 14 PCs as selected by optim.a.score.

**Supplementary 3.**
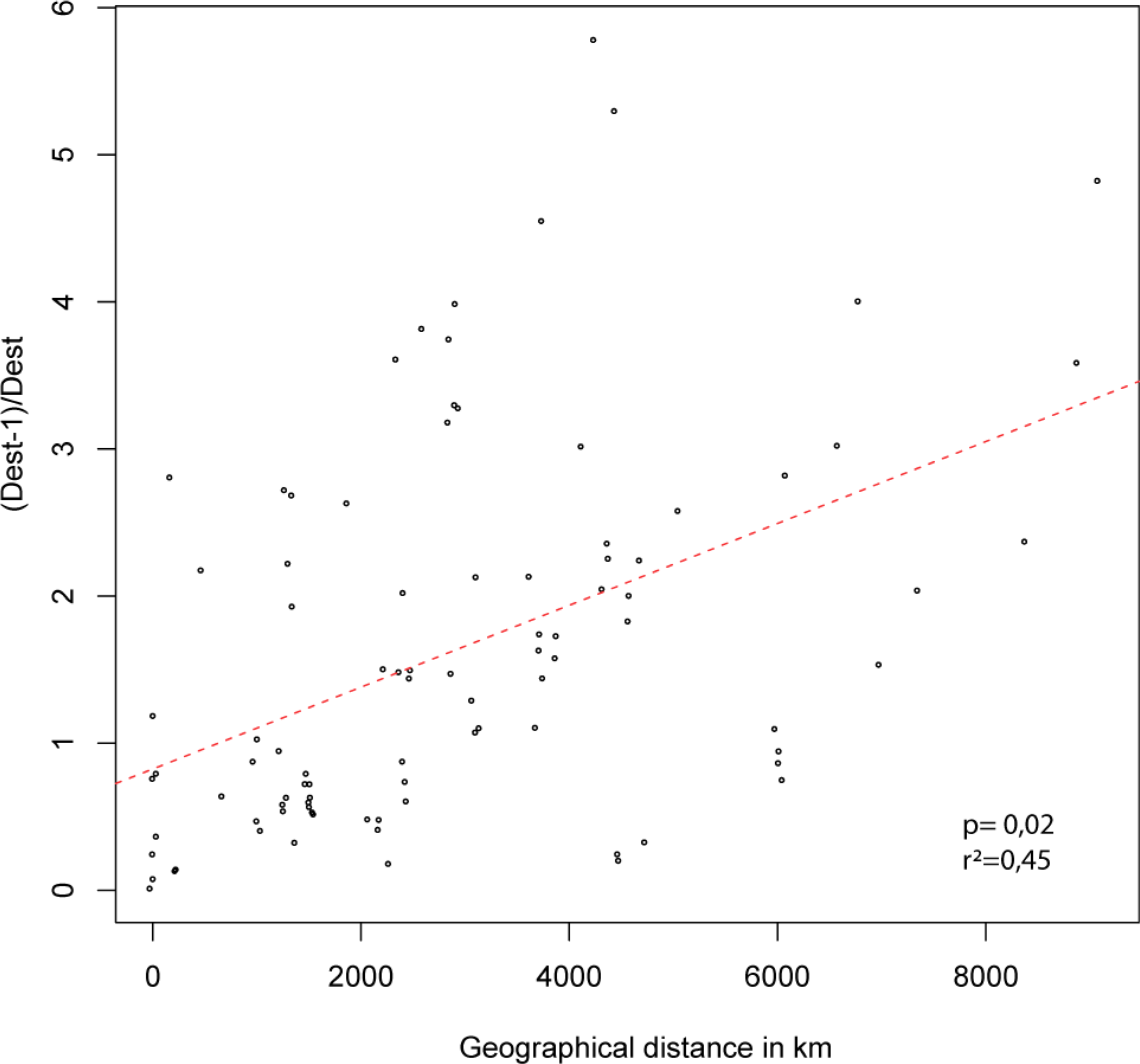
IBD for all European populations, including Punta del Hidalgo. The corresponding p-value for the Mantel-test and the correlation coefficient of the regression line r² are indicated bottom right. Linearized Jost Dest is projected against geographical distance in km.

**Supplementary 4a.**
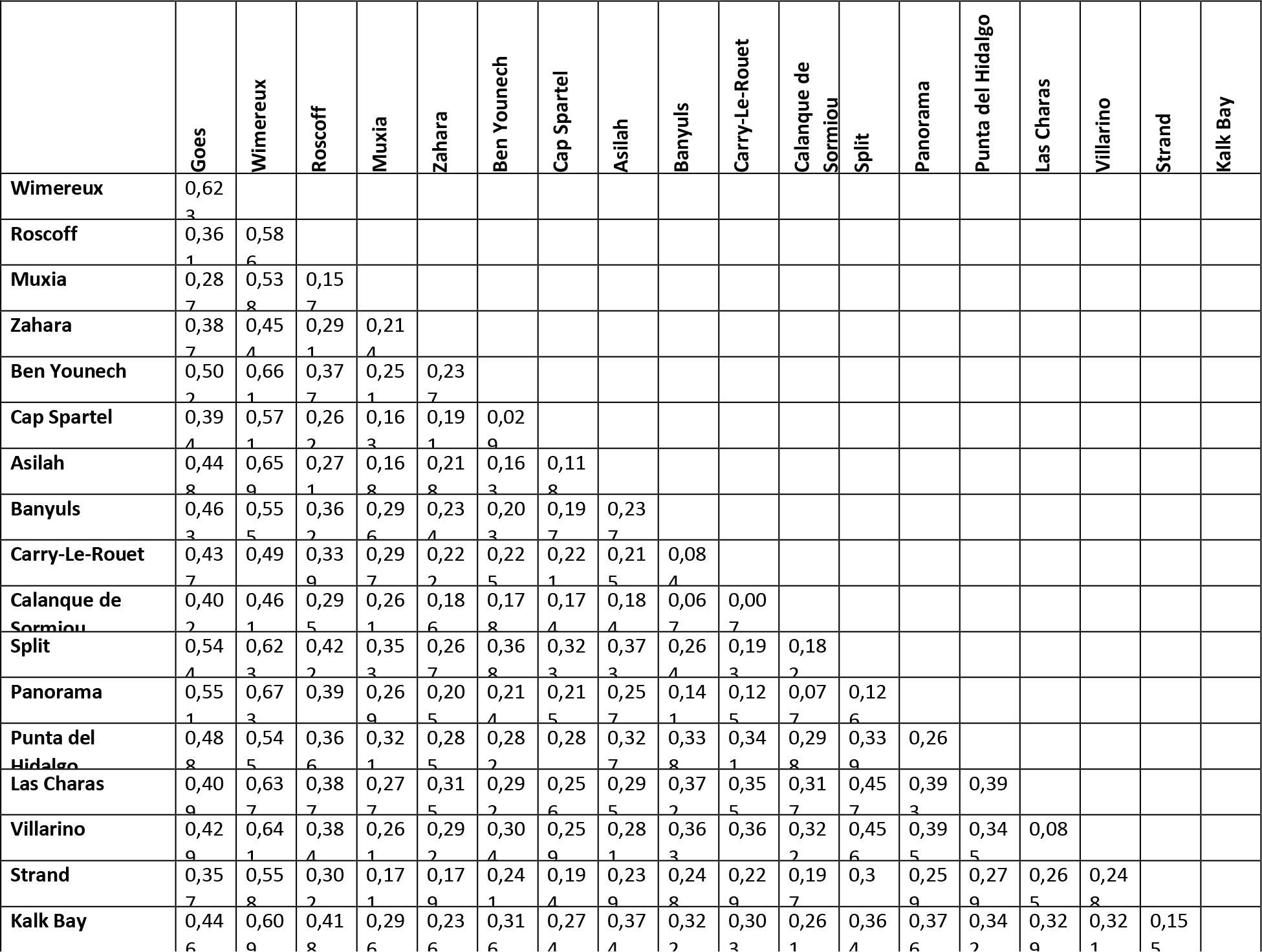
Pairwise Fst for all of the populations.

**Supplementary 4b.**
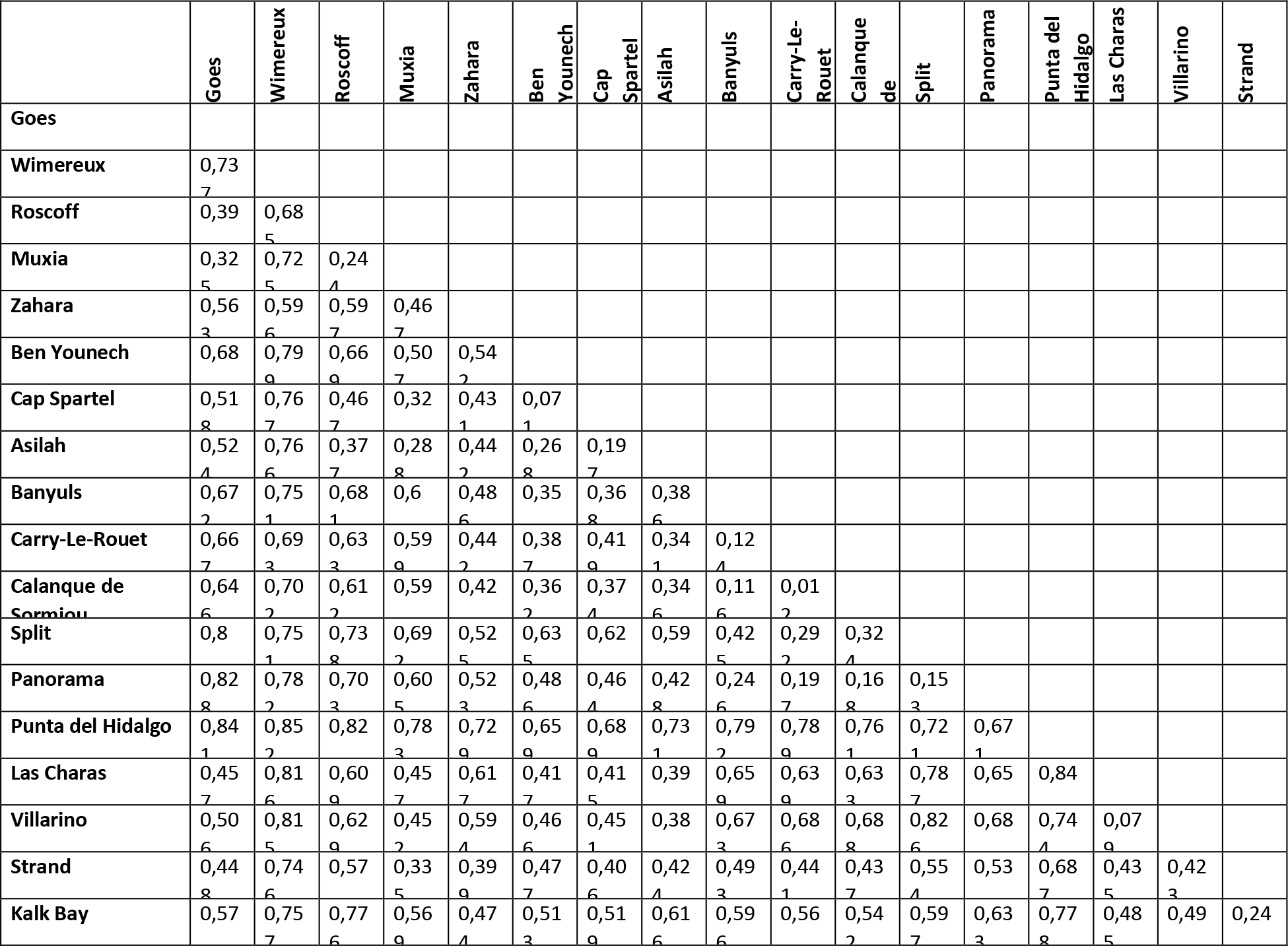
Pairwise population differentiation as measured by Jost linearized D.

